# Life history effects on the molecular clock of autosomes and sex chromosomes

**DOI:** 10.1101/024281

**Authors:** Guy Amster, Guy Sella

## Abstract

One of the foundational results of molecular evolution is that the rate at which neutral substitutions accumulate on a lineage equals the rate at which mutations arise. Traits that affect rates of mutation therefore also affect the phylogenetic “molecular clock”. We consider the effects of sex-specific generation times and mutation rates in species with two sexes. In particular, we focus on the effects that the age of onset of male puberty and rates of spermatogenesis have likely had in extant hominines (i.e., human, chimpanzee and gorilla), considering a model that approximates features of the mutational process in most mammals and birds and some other vertebrates. As we show, this model helps explain and reconcile a number of seemingly puzzling observations. In hominines, it can explain the puzzlingly low X-to-autosome ratios of substitution rates and how the ratios and rates of autosomal substitutions differ among lineages. Importantly, it suggests how to translate pedigree-based estimates of human mutation rates into split times among apes, given sex-specific life histories. In so doing, it helps bridge the gap between estimates of split times of apes based on fossil and molecular evidence. Finally, considering these effects can help to reconcile recent evidence that changes in generation times should have small effects on mutation rates in humans with classic studies suggesting that they have had major effects on rates of evolution in the mammalian phylogeny.

## Introduction

Most of our inferences about species split times on short phylogenetic time scales rely on the neutral molecular clock. According to the Neutral Theory, the number of substitutions *K* that accumulate in a lineage over *T* years (e.g. since the split from another species) is 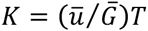 where ū and 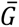 are the average mutation rate per generation and generation time, respectively (1). To infer split times, estimates of the yearly mutation rate 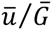 are therefore required for the lineage in question. Estimates of yearly rates generally derive from securely dated fossils on other lineages or from measurements of mutation rates in extant species (2-5). Using these estimates for dating thus necessitates an understanding of the way that yearly mutation rates may change over time.

Neutral substitution patterns in mammals offer some insights. Variation in yearly mutation rates on phylogenetic time scales can be assessed by comparing the number of neutral substitutions along two branches leading from a common ancestor to extant species. These comparisons show marked variation in yearly rates on autosomes. For example, there are 50% fewer substitutions on the human branch compared with rodents (6) and 26% fewer compared to baboons, with more moderate differences among lineages of great apes (6-9). The average yearly rates are also negatively correlated with generation times (and their correlates) in extant mammals, leading to the notion of a generation time effect on the molecular clock (6, 10, 11).

Neutral substitutions rates vary not only among taxa but also between sex chromosomes and autosomes. For brevity, we consider the relative rates on X and autosomes, but these considerations extend naturally to Y (or ZW). Because autosomes spend the same number of generations in both sexes while the X spends twice as many generations in females, rates of neutral substitutions on autosomes reflect a greater relative contribution of male mutations than on the X. In a wide range of taxa, neutral substitutions rates on autosomes are greater than on the X (or lower than on the Z) thus revealing a male biased contribution to yearly mutation rates (12). Moreover, observed X-to-autosome ratios are extremely variable, ranging between 0.76-0.9 in great apes and up to 1.0 in surveyed mammals, suggesting that the degree of male bias itself greatly varies on phylogenetic time scales (12, 13).

Our current understanding of mutation may help tie these observations together (5). Pedigree studies in humans and chimpanzees now establish that most mutations are paternal and that the paternal but not the maternal contribution increases strongly with age (4, 14-16). This has long been thought to be true because germ-cell division is arrested before birth in females but proceeds continuously post-puberty in males (5, 17-20). The same reasoning may extend to most mammals and birds and other vertebrate taxa in which oogenesis ceases at birth or hatching (21-23). These considerations suggest that maternal and paternal generation times would affect the rate of the molecular clock differently. They also suggest that in males, age of puberty and the rate and architecture of germ cell division would be important (5). Given the variation among closely related species, we know that life history traits and the process of spermatogenesis changed over phylogenetic time scales. Here we ask how such changes would affect the molecular clock on X and autosomes.

## Model

### The molecular clock with two sexes

We model the accumulation of neutral substitutions allowing for different generation times and mutation rates in males and females. The expected number of substitutions *K* over *T* years can be expressed in terms of the expectations for the male and female generation times (defined as their average age at parturition), *G*_*M*_ and *G*_*F*_, and mutation rates per generation, *μ*_*M*_ and *μ*_*F*_. On autosomes, it is

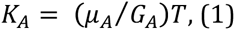

where 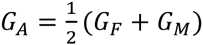 and 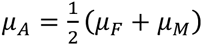 are the expected sex-averaged generation time and mutation rate per-generation on an autosomal lineage (cf. SI Section 1 for rigorous derivations). This relationship can be viewed as the expected number of generations (*T*/*G*_*A*_) times the expected mutation rate per generation (*μ*_*A*_) or, equivalently, as the expected mutation rate per year (*μ*_*A*_/*G*_*A*_) times the number of years (*T*). By the same token, on the X

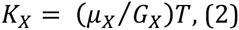

where in this case 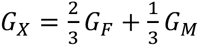 and 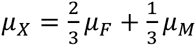 are the expected sex-averaged generation time and mutation rate per-generation on a X lineage.

The X-to-autosome ratio of the number of substitutions follows. In terms of the ratios of male-to-female generation times, *G*_*M*_/*G*_*F*_, and mutation rates, *α* = (*μ*_*M*_/ *μ*_*F*_ (also referred to as the male mutation bias), it takes the form

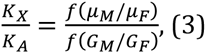

where 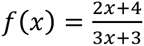.

### Sex and age dependent mutation rates in hominines

We model male and female mutation rates per generation, *μ*_*M*_ and *μ*_*F*_ based on the current understanding of the process of accumulation of germline mutations in great apes (5, 18). Notably, we assume that mutations accumulate linearly with the number of germ-cell divisions, where the rate per division varies at different stages of development (5). This is a natural assumption for replicative mutations and has recently been suggested to apply to non-replicative mutations as well, if efficiently repaired ((24); cf. Discussion for other kinds of mutations).

In females, all oogonial mitotic divisions occur before birth, so the number of mutations should have no dependence on the age of reproduction ((22), but see (25, 26)). This assumption is supported by pedigree studies in humans and chimpanzees finding that maternal age has little or no effect on mutation rates (4, 14-16). We therefore model the female per-generation mutation rate (*μ*_*F*_) as a constant.

Germ-cell divisions in males exhibit two main phases: pre-puberty, starting from the zygote through the proliferation of germ cells in the growing testis, and post-puberty, with continuous divisions in the adult testis during spermatogenesis. While testis mass vary considerably among great apes (13), the number of cell divisions should increase only logarithmically with mass. We therefore expect it to have a small effect and approximate the expected number of mutations pre-puberty as constant (*C*_*M*_). Post-puberty, one germ cell division is thought to occur at each seminiferous epithelial cycle (27), where the length of the cycle (τ) varies among great apes (Table 1). We therefore model the number of mutations accumulated between the age of puberty *a*_*P*_ and reproduction *a*_*R*_ as (*a*_*R*_– *a*_*P*_) (*D*_*M*_/ τ) where *D*_*M*_ is the number of mutations per spermatogenic division. This form is consistent with the apparent linear increase in mutation rates with paternal age observed in pedigree studies (4, 14-16). Taken together and averaging over the distribution of puberty and reproductive ages in the population (with means *P* and *G*_*M*_*-I*, where *G*_*M*_ is the average age at offspring birth and *I* is the gestation time), we model the average mutation rate per generation in males by

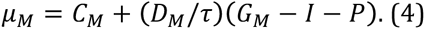

**Table 1:** Estimates of spermatogenesis and life history parameters and summaries of neutral substitution patterns in hominines. More details about various parameter estimates are provided in SI Section 4.

|  | Spermato-<br>genesis rate<br>(days) | Age of puberty<br>in males<br>(years) | Populations | Maternal gen.<br>time (years) | Paternal gen.<br>time (years) | Average gen.<br>time (years) | Ratio of male<br>to female gen.<br>time | Ratio of X to<br>autosome<br>divergence | Autosomal<br>branch length<br>(relative to the<br>human branch) |
| --- | --- | --- | --- | --- | --- | --- | --- | --- | --- |
| Human | 16<sup>(28)</sup> | 13<sup>(29)</sup> | Hunter-gatherers<br>Modern<br>countries | 26.9*<br>27.3<sup>(30)</sup> | 33.8*<br>30.8<sup>(30)</sup> | 30.4<br>29.1 | 1.26<br>1.13 | 0.8<sup>(13)</sup> / 0.79<sup>(13)</sup> | - |
| Chimpanzee | 14<sup>(31)</sup> | 7.5<sup>(32)</sup> | Western<br>Eastern<br>Weighted<br>average | 26.3<sup>(33)</sup><br>24.8<sup>(33)</sup><br>25.2<sup>(33)</sup> | 24.3<sup>(33)</sup><br>24<sup>(33)</sup><br>24.1<sup>(33)</sup> | 25.3<br>24.4<br>24.6 | 0.93<br>0.97<br>0.96 | 0.76<sup>(13)</sup> /<br>0.79<sup>(13)</sup> | +0.4%<sup>(7)</sup> /<br>+0.7%<sup>(9)</sup> /<br>+2.9%<sup>(34)</sup> |
| Gorilla | - | 7<sup>(35)</sup> | Mountain<br>gorillas | 18.2<sup>(33)</sup> | 20.4<sup>(33)</sup> | 19.3 | 1.12 | 0.9<sup>(13)</sup> /<br>0.88<sup>(13)</sup> | +6.1%<sup>(7)</sup> /<br>+8.7%<sup>(9)</sup> /<br>+11%<sup>(34)</sup> |

\* Revised from (30); see SI Section 4.

A similar model may provide a reasonable approximation for vertebrate species in which oogenesis ceases at birth or hatching (e.g. most eutherian mammals, monotremes, birds, elasmobranchs, cyclostomes and a few teleosts (22, 23), but see (25, 26)), with some modifications (e.g., accounting for intermittent spermatogenesis (21)).

In general, incorporating a mutational model into the model of the molecular clock entails averaging over the distribution of mutational parameters both within a population at a given time and over phylogenetic time scales. Notably, this averaging should account for the way the rate of the molecular clock depends jointly on generation times and mutation rates per generation in males and females. In SI Sections 1 and 2, we describe how to perform the averaging in general (e.g., nonlinear) mutational models and justify its form (i.e., Equation 4) for our illustrative case. Specifically we show that the expected number of substitutions is insensitive to the variance of the number of mutations or to the variance of the age at reproduction, thus justifying our considering only the means.

### Parameter values and ranges

To study how changes in life history traits and spermatogenesis in hominines would have affected the rate of the molecular clock, we would like to assign realistic values to the parameters of the mutational model. Lacking evidence to the contrary, we assume that the parameters associated with the rates of mutation per germ-cell division at different developmental stages remained constant throughout the great ape phylogeny. We infer these parameters from the relationship between mutation rates and paternal ages in the largest human pedigree study published to date (4), which yields: *C*_*M*_ = 6.13×10^−9^, *D*_*M*_ = 3.33×10^−11^ and *μ*_*F*_ = 5.42×10^−9^ per base pair (see SI Section 4 for details). In contrast, life history and spermatogenesis parameters, *G*_*M*_, *G*_*F*_, *P* and τ, are known to vary among extant great apes and Old World Monkeys (OWM), and even among populations of individual species (cf. Tables 1 and S3-7 and references therein), indicating that they have changed throughout the phylogeny. We use the variation among extant species to guide our choice of plausible ranges for these parameters. The parameter values based on our survey of the literature, including life history, physiological and mutational studies, are summarized in Table 1 (see SI Section 4 for further details).

There is considerable uncertainty about the values of these parameters in extant species, let alone about their variation on the hominine phylogeny. For example, estimates often vary among populations of extant species (e.g., maternal generation times in communities of western chimpanzees; cf. Table S5) and are based on different methodologies and operational definitions (e.g., puberty age in males is measured by testicular descent, testicular enlargement, hormonal changes or spermatozoa production) (see SI Section 4 for more details). Moreover, to the best of our knowledge, only subsets of the parameters were measured in most species. Lastly, extant species afford only a small sample of life history and spermatogenesis parameters values, so these values were likely to vary over larger ranges within the phylogeny. For all these reasons, the parameter ranges estimated from extant species should be treated only as a rough guide for their ranges over the phylogeny.

## Results

### The autosomal molecular clock

We first consider the effects of male and female generation times. To this end, we distinguish between the effects of the ratio of male-to-female generation times and of the sex-averaged generation time. In general, the impact of changing the ratio or average of generation times will depend on the way mutation rates vary with sex and age (see Discussion and SI Section 3 for more details). If, for example, mutation rates increase more rapidly with paternal than maternal age, then increasing the ratio of male to female generation times necessarily increases the mutation rate per year.

Predicting the effect of increasing the average generation time requires additional assumptions. As an example, we assume that females and males contribute a constant number of mutations pre-puberty and that mutations in males post puberty accumulate at a constant rate (i.e., the functional form but not the parameter values of our hominine mutational model). If we further assume that the proportion of the male generation time spent pre- and post-puberty remains constant (36, figure S1), then increasing the average generation time will decrease the rate of the molecular clock, consistent with what is seen in phylogenetic studies (6, 10, 11). If, instead, we assume that puberty age remains constant while the average generation time increases, which better captures the variation observed in extant hominines (cf. Table 1 and S1) and is therefore what we assume below, then the effect on the molecular clock can go either way and depends on parameter values (i.e., on *C*_*M*_, *D*_*M*_ and *μ*_*F*_; cf. SI Section 3 and (5, 24)). We return to the generation time effect in broader phylogenetic contexts in the Discussion.

Here we focus on our model and parameter estimates for hominines, which strongly suggests that changes to the ratio of generation times have a much greater potential impact than changes to the average generation time (Figure 1A). Based on our mutation model, replacing the standard (often implicit) assumption of equal generation times in males and females with estimates of their ratio in current hunter-gatherer populations increases yearly mutation rates by ∼10%. Further considering that the ratios in extant hominine species vary between 0.92 for western chimpanzees and 1.26 for hunter-gatherers (cf. SI Section 4) suggests that yearly mutation rates could be anything between 4% lower to 10% higher than obtained assuming a ratio of 1 (unless noted otherwise, when we vary a single parameter, other parameters are assigned human estimates). By comparison, varying the average generation time between estimated values of 19, for gorillas, and 30.4, for extant hunter-gatherer populations, impacts rates by less than 2% (see also (24)).

**Figure 1:**
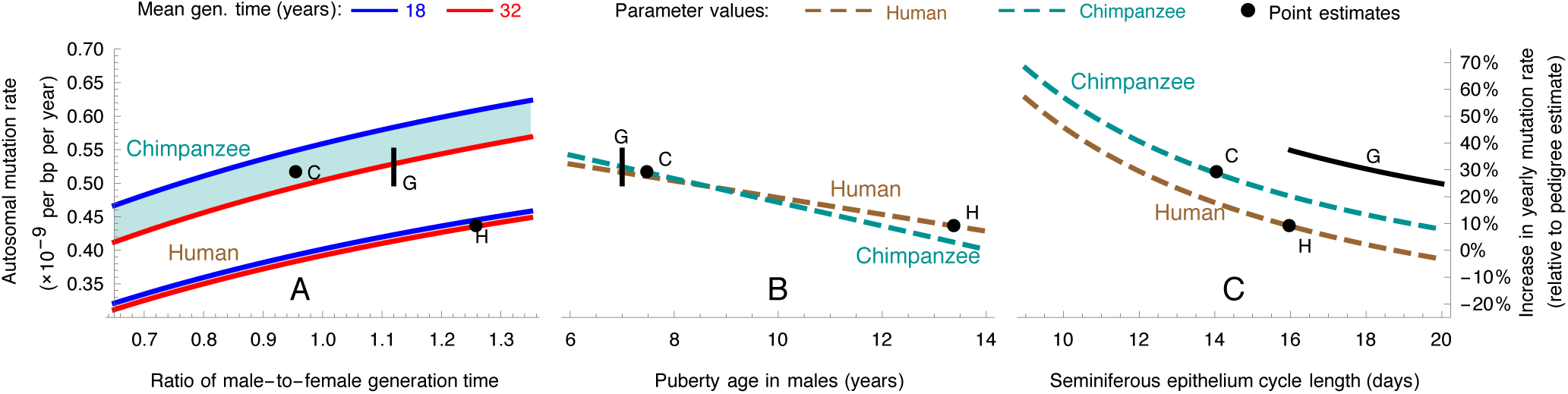
Predicted yearly mutation rates on autosomes as a function of the ratio of male to female generation times (A), male age of puberty (B) and rate of spermatogenesis (C). Rates are measured relative to the estimate of 0.4 · 10^−9^ per bp per year reported in Kong et al. (4), which is based on the same data that we used to fit the parameters of our mutational model (cf. Model section). In each panel, we vary one parameter, while fixing other parameters to their estimated values in extant humans (brown) or chimpanzees (green) (see Table 1). Point estimates for the rates in humans and chimpanzees are shown with black points. Note that these estimates do not coincide with those reported in pedigree studies, because ours account for the predicted effects of life history and spermatogenesis parameters. Estimates for gorillas, where the rate of spermatogenesis has not been measured, are marked by a black line corresponding to a range of spermatogenesis rates between 16 and 20 days.

Male puberty age and rate of spermatogenesis also affect yearly mutation rates (Figure 1B and C; (5, 24)). Notably, both an earlier onset of puberty and an increased rate of spermatogenesis increase the yearly mutation rates in males, resulting in an increased rate on autosomes. Varying their values within the range known for hominines markedly affects yearly rates, by ∼18% for the male puberty age and ∼8% for the rate of spermatogenesis.

Considered jointly, we would expect these factors to generate differences in the branch lengths leading to extant hominines. Assuming parameter estimates for extant humans and chimpanzees, for instance, our model suggests that the human branch would be 15% shorter. If instead we make the more plausible assumption that the yearly mutation rate in the ancestor was between extant estimates and that the rates on each lineage changed gradually, we would still expect the human branch to be somewhat shorter. The human branch has been estimated to be 0.4%-2.9% shorter (7, 9, 34); these estimates include the contribution of ancestral polymorphism in common to both branches, suggesting that the difference between the human and chimpanzee specific branches is larger.

The lack of estimates for the rate of spermatogenesis in gorillas prevents us from making similar predictions for their relative branch lengths. Nonetheless, rates of spermatogenesis and relative testis mass are positively correlated in mammals, and the variation has been attributed to differences in the intensity of sperm competition in different mating systems (e.g., monogamous, polygamous or multi-male and multi-female). In agreement with this hypothesis, the relative testis mass and rate of spermatogenesis in chimpanzees (0.27% of body weight and 14 days), which have a promiscuous mating system, are greater than in humans (0.07% of body weight and 16 days), which are largely monogamous, where the relative testis mass in polygynous gorillas (one male controls reproductive access to many females) are smaller than in both (0.02% of body weight) (13, 37). This would suggest that the rate of spermatogenesis in gorillas is lower than in humans. Varying the length of the spermatogenetic cycle between 16 days (its value for humans) and 20 days yields predicted branch lengths 14% to 26% longer than the human branch, respectively (and between 3% shorter to 6.5% longer than the chimpanzee branch). These predictions accord with current estimates suggesting that the gorilla branch is between 6.1%-11% longer than the human branch (again, without accounting for the ancestral contribution) (7, 9, 34).

Lastly, we consider the effects on estimates of split times between humans and chimpanzees. Usually these estimates derive from dividing the neutral divergence on the human lineage (∼0.8% per bp, after subtracting the contribution of polymorphisms in the ancestral population; see (38) and Table S9) by estimates of the yearly mutation rate (∼0.4×10^−9^ per bp per year, from the Kong et al. pedigree study (4)), suggesting a split time of ∼10 MYA ((39) and SI Section 5). Pedigree-based estimates, however, do not account for most of the effects that we have considered (cf. Discussion). For instance, replacing the standard (implicit) assumption of equal generation times in males and females with the ratio estimated in extant hunter-gatherers increases yearly rates and shortens split time estimates (Figure 1). Nonetheless, the extensive variation in the ratio among extant populations (cf. Tables 1 and S3) suggests that it could have evolved rapidly on the human lineage, introducing considerable uncertainty. We are also uncertain about changes to the age of male puberty and rate of spermatogenesis. The earlier age of puberty in chimpanzees and evidence for earlier puberty in *Homo erectus* (e.g., from Nariokotome boy dated ∼1.5 MYA (40, 41)) suggest that the onset of puberty could have occurred at younger ages during most of the human lineage. The rate of spermatogenesis is higher in chimpanzees, suggesting that it could have been higher on the human lineage. While it is difficult to translate these lines of evidence into estimates of split times, it is interesting to note that they all suggest that the split time was more recent than the naïve pedigree-based estimates would suggest (i.e., ignoring these factors). For example, if the average yearly rate on the human lineage was between the rates in extant humans and chimpanzees then our model suggests a split time between 7.7-9.1 MYA (Figure 1A and SI Section 5). If we further allow individual life history parameters to range between their values in extant humans and chimpanzees then the lower bound on the split time is further reduced to ∼6.6 MYA (see Figure 4).

## The relative rates of the molecular clock on X and autosomes

Differences in neutral rates of substitutions on sex chromosomes and autosome have generally been attributed to differences in mutation rates among sexes. In fact, their ratios have been widely used to infer the ratio of male-to-female mutation rates, α = *μ*_*M*_/*μ*_*F*_, in lineages from mammals, birds, flies, fish and plants (10, 12, 42-51). The inferences from X-to-autosome ratios rely on Miyata’s formula (52)

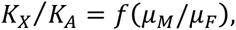

where 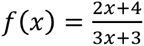 (For brevity, we focus on X and autosome, where the modifications for other sex chromosomes are straightforward; SI Section 1). Critically, Miyata’s formula assumes equal male and female generation times. When this assumption is relaxed, the X-to-autosome ratio also depends on the ratio of male-to-female generation time:

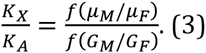

This suggests a theoretical range between 1/2 and 2 compared to a range between 2/3 and 4/3 based on Miyata’s formula.

Ignoring differences in generation times could introduce substantial biases in estimates of the male mutation bias, *α* (Figure 2). Notably, when *α* is large, an incremental increase in *α* has a smaller effect on the X-to-autosome ratio (because of the form of *f*; Figure 2A). In this case, even moderate differences in the ratio of generation times translate into large biases in estimates of the mutational bias (Figures 2B and S2). For example, if the mutational bias is *α*=3.9 (the inferred value for hunter-gatherers based on our model and accounting for their generation times) and the difference in generation times between sexes is between 25% greater in males and 25% greater in females, then ignoring the difference in generation times induces a bias in estimates of *α* of -23% to 46%. Hence, as differences in generation times among sexes are common (33, 53), it is likely that many current estimates of *α* suffer from a substantial bias (Figure 2C). This suggests that information about differences among sexes in generation times (e.g., from extant species) is key to assessing the uncertainty in inferences about mutational biases.

**Figure 2:**
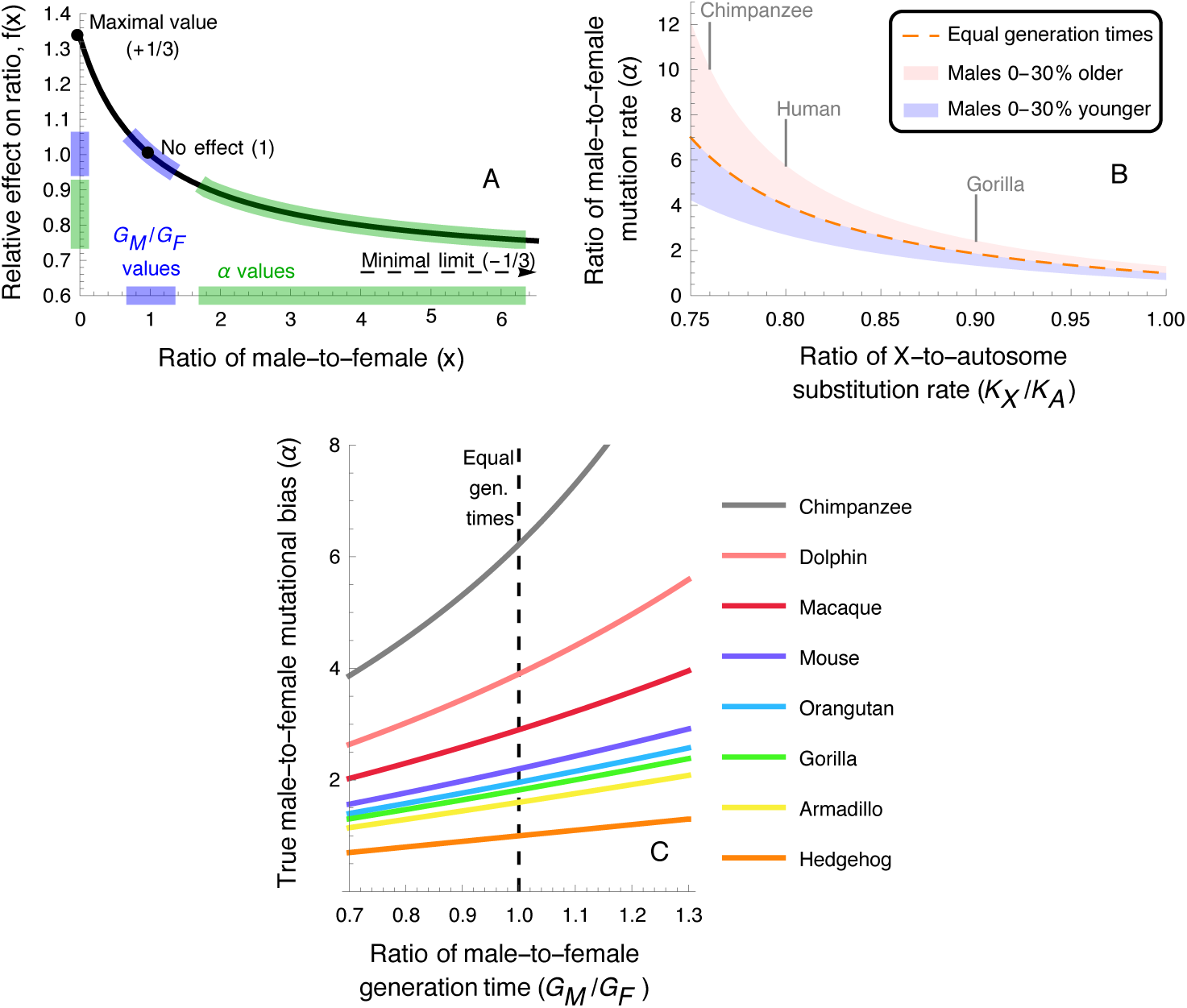
The effects of the male-to-female ratio of generation times on inferences of male biased mutation. (A) The function *f* mediating the effects of ratio of male-to-female generation times and mutation rates on the X-to-autosome ratio of substitution rate. The effect (relative to a ratio of 1) is bound between -1/3 and 1/3. The highlighted regions correspond to the ranges of male-to-female ratios of generation time (blue) and of mutation rate (green) observed in hominines. (B) The effects of mild differences between male and female generation times on inferences of the ratio of male-to-female mutation rate (*α*) based on Miyata’s formula. (C) The potential biases in estimates of the ratio of male-to-female mutation rates in 8 mammalian species, as a function of the ratio of male-to-female generation times. The curve for each species is based on the X-to-autosome divergence ratio reported along its lineage (12, 13) (neglecting additional effects, e.g. differences in the ancestral contributions and in base composition between X and autosome). The dashed black line marks the estimates based on Miyata’s formula, which assumes equal generation times in males and females.

## X-to-Autosomes ratios in hominines

Our model for hominines relates mutation rates with the underlying life history and spermatogenesis parameters, allowing us to investigate how changes to these parameters would affect X-to autosome ratios. We first consider the effect of female and male generation times. Increasing the paternal generation time has opposing effects on the ratio, as it increases both the male mutation bias (*α*) and the ratio of generation times (*G*_*M*_/*G*_*F*_). In contrast, changing the maternal generation time affects only the ratio of generation times. As a result, changing the maternal generation time has a greater effect on the X-to-autosomes ratio (Figure 3A). For example, varying maternal age between 18 and 25 (roughly the estimated range in hominines) decreases the ratio by ∼6%, whereas varying paternal age within this range decreases the ratio only by ∼2.5%. In turn, both earlier male puberty and an increased rate of spermatogenesis increase mutation rates in males resulting in a lower X-to-autosome ratio (Figure 3B and C). Varying these parameters within the known ranges for hominines and assuming a spermatogenesis cycle of 18 days for gorillas yields effects of 2.8% for male puberty and 2.4% for spermatogenesis (smaller than the effect of maternal age).

**Figure 3:**
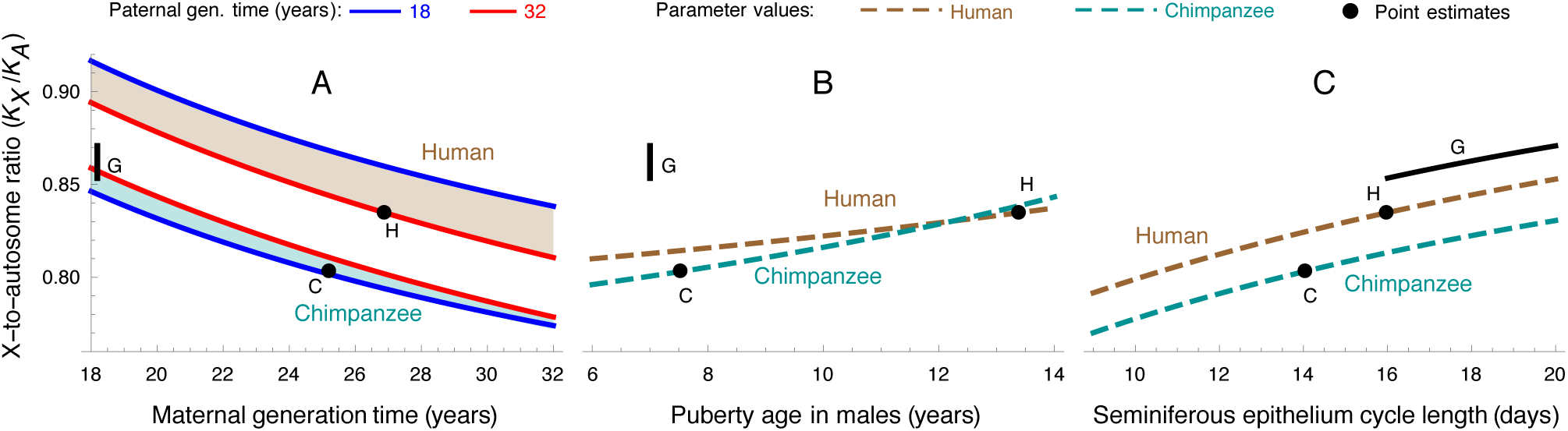
Predicted X-to-autosome ratios of substitution rates as a function of male and female generation times (A), male age of puberty (B) and rate of spermatogenesis (C). In each panel, we consider the effects of one of the parameters while the others are fixed for their estimates in extant humans (brown) or chimpanzees (green) (see Table 1). Point estimates for the ratio in humans and chimpanzees are marked by black points. Estimates for gorillas, where the rate of spermatogenesis has not been measured, are marked by a black line corresponding to a range of spermatogenesis rates between 16 and 20 days.

In hominines, X-to-autosome ratios have garnered considerable attention, in part because of what seems like inexplicably low ratios (9, 13, 54). Estimates of the ratios of neutral divergence are ∼0.76 for the chimpanzee lineage, ∼0.8 for the human lineage and ∼0.9 for gorillas (13). These estimates, however, do not correspond directly to the ratios that we are considering. Notably they include the ancestral contributions, which are likely greater on autosomes (9, 54, 55), suggesting that the ratio after the species splits would be closer to 1. Also, they do not control for differences in base composition between X and autosomes. In that regard, we note that the standard practice of dividing by divergence to an outgroup only complicates the interpretation by compounding estimates of the X-to-autosome ratios in hominines with the ratio on the outgroup lineage. Finally, existing estimates do not account for the asymmetry in branch lengths. For all these reasons, we consider the current estimates to be rough.

These caveats notwithstanding, we can ask whether the observed values could be explained in terms of our mutational model. Using parameter estimates from extant species, we would predict an X-to-autosome ratio of 0.80 on the chimpanzee lineage and 0.83 on the human lineage. We further assume that weaker sperm competition yielded lower rates of spermatogenesis in gorillas. Varying the length of the spermatogenetic cycle between 16 (its value for humans) and 20 days yields ratio predictions between 0.85 and 0.87, correspondingly. Taken together these estimates recover the ordering of ratios among lineages as well as the rough magnitude of the reduction below 1. Further considering the ranges in which individual parameters vary in extant species can explain both a smaller ratio in chimpanzees and a greater difference between the chimpanzee and gorilla ratios (Table S8). Thus, it is plausible that observed X-to-autosome ratios could be explained by the effects of life history (e.g., generation times and male puberty age) and rates of spermatogenesis, without recourse to more elaborate scenarios.

Lastly, we consider the relative contributions of rates of spermatogenesis and life history traits. Presgraves and Yi (13) pointed out that variation in mutation rates rather than complex speciation (9) could explain the X-autosome ratios in great apes, suggesting that differences in the intensity of sperm competition might be responsible. Assuming that testis mass has a negligible effect on yearly mutation rates, this is equivalent to attributing the difference in ratios to variation in rates of spermatogenesis. In our model, this explanation would require the seminiferous epithelial cycle length in gorillas to be at least 65 days long, exceeding the maximal value measured in mammals by almost fourfold (Figure S3). While we cannot rule this out (let alone differences in the architecture of cell division in spermatogenesis; cf. Discussion), our mutational model alongside other parameters estimates in extant species would suggest that life history traits, particularly differences in maternal generation times (Figure 1A) and male puberty age (Figure 1B), were likely to have contributed more.

## Discussion

Human pedigree studies now allow for direct measurements of mutation rates, and should therefore allow for more accurate and precise dating of species split times and other evolutionary events (e.g., Out-of-Africa migrations). Doing so, however, requires mutation rates from pedigrees to be translated into yearly substitution rates on a phylogeny. The standard approach is to divide the sex averaged mutation rate per generation (in a study) by estimates of the sex averaged generation time along the lineage under consideration. This calculation implicitly assumes that the mutation rate per generation remains constant, reflecting an extreme interpretation of the generation time effect (5) (which, as we discuss below might be justified for mammals with short lifespans). Importantly, it also ignores differences in generation times among sexes. Yet, our results show that such differences have a substantial effect on yearly rates, in fact a much greater effect than changes in the sex-averaged generation time. Moreover, because paternal rates strongly depend on age, differences in paternal age between a given pedigree study and the evolutionary lineage in question could also have a substantial effect on estimates of yearly rates (5). Thus, translating the results of pedigree studies into yearly mutation rates is not as straightforward as it seems, and requires consideration of these life history factors.

A more sensible approach is to fit the results of pedigree studies with a mutational model that reflects the dependency of yearly rates on age and sex. This allows yearly rates along a lineage to be inferred based on estimates of female and male generation times and measures of uncertainty to incorporate information about variation in generation times and other factors (e.g., male puberty age and rate of spermatogenesis) in extant populations and related species. Our simple mutational model for hominines illustrates such an approach and leads to substantial changes in estimates of yearly rates and corresponding split times. Most notably, it revises split times downwards, e.g., from ∼10MYA to as low as ∼6.6 MYA for the human-chimpanzee split (see Figure 4 and Table S8).

While we would argue that our mutational model offers a considerable improvement, it may well be only a rough approximation of reality. For example, molecular evolutionary patterns suggest that rates of certain types of mutations (e.g., CpG transitions; (56, 57)) may depend on absolute time in addition to their dependency on the number of germ-cell divisions at different developmental stages (5). This would also suggest that maternal age should affect the rates of certain types of mutations (5). Lastly, there is still uncertainty about the architecture of germ-cell division during spermatogenesis. It has been suggested that *A*_*pale*_cells - the stem cells from which sperm are generated - are replenished by, or experience turnover with *A*_*dark*_spermatogonial stem cells throughout adulthood, resulting in fewer germ-cell divisions between puberty and reproduction (27, 58, 59). Revising the estimates for the number of cell divisions would help reconcile the mutation rate per-spermatogenic cell division (*D*_*M*_) with the per-division rates in females, in males pre-puberty (5) and in other species (e.g., our inferred *D*_*M*_ is 2-20 times lower than in *E. coli, S. cerevisiae, C. elegans, A. thaliana, D. melanogaster* and *M. musculus*; cf. Table S2 and (60, 61)). In terms of the hominine mutational model, such a revision would introduce an additional parameter for turnover allowing for greater variation among species in male mutation rates post-puberty (59). As pedigree studies increase in sample size, their methodology improves (e.g. in accounting for false negatives; (5)) and they are performed in more species, we will surely learn more about the dependence of different types of mutation on sex and age, allowing for refinements of our mutational models.

These reservations notwithstanding, our simplified mutational model already suggests some qualitative predictions that appear to be consistent with observation. For instance, our predicted mutation rates in chimpanzees fall within confidence intervals of a recent pedigree study by Venn and colleagues (15). We also predict the reduction in branch length leading to humans compared to gorillas; although a quantitative comparison awaits better estimates (e.g., accounting for the ancestral contributions).

In addition, our simplified model provides sensible qualitative predictions for the X-to-autosome ratios in hominines. Notably, it predicts that the ratios should be substantially below 1 and lower on the human and chimpanzee lineages than on that of gorilla. Once again, quantitative comparisons will require ancestral contributions to be taken into account and controls for base composition. We note that while differences in life history and spermatogenesis could explain variation in X-to-autosome ratios, it cannot explain the variation in human-chimpanzee divergence along the X chromosome, which has been suggested to reflect the footprints of linked selection in the ancestral population or during speciation in the presence of gene flow (9, 54).

Our results also bear on the recently invigorated discussion about split times in the great ape phylogeny (39). Recent pedigree studies suggest that yearly mutation rates in humans are more than twofold smaller than previous estimates based on the fossil record. Assuming that pedigree-based estimates reflect yearly rates along the human lineage suggests more than a twofold increase in split time estimates, placing the human-chimpanzee split at ∼10 MYA instead of ∼4 MYA, based on fossil calibration for the human-orangutan or human-OWM splits (cf. SI Section 5 and (38, 39)). Attempts to reconcile pedigree and fossil based estimates appear to be moving from both ends. From one end, it has been suggested that mutation rates experienced a slow down in the great ape phylogeny (39). From the other, the new rate estimates from pedigrees triggered a reexamination of the veracity of evidence for the phylogenetic positioning of ape and primate fossils (62). In particular, it has been argued that upper bounds (i.e., the estimates that are farthest back in time) used for the human-orangutan and human-OWM split times are too conservative and that they should extend as far back as 23 and 34 MYA (62); these bounds are consistent with mutation rates as low as 0.65 and 0.84 10^−9^ per bp per year, respectively (cf. SI Section 5). More realistic models of the molecular clock can inform this discussion by suggesting plausible ranges for changes to yearly mutation rates, better delineating both the extent of a mutational slow down and the fossil positions that would in fact contradict pedigree based estimates.

As an illustration, we use our mutational model alongside estimates of life history and spermatogenesis parameters from extant species to suggest plausible ranges for yearly mutation rates on the great apes phylogeny (Figure 4; Table S8 and SI Section 5). Specifically, we allow each of the parameters on branches following a split to vary independently within the range observed in extant populations descended from that split. For the parameter estimates lacking in orangutans and gorillas, we assume intermediate values between extant OWM and extant humans and chimpanzees. This implies that the ranges for individual parameters and for the resulting yearly mutation rates become larger when we go farther back in time. Interestingly, we find that the ranges expand only upwards, supporting the notion of a mutational slow down. Moreover, the inferred ranges could, in theory, reconcile the apparent discrepancy between yearly rates of 0.4×10^−9^ per bp observed in pedigree studies, and yearly rates between *0*.6 - 1.0×10^−9^ per bp inferred from the fossil record (cf. SI section 5 for more details). We note, however, that assuming that life history and spermatogenesis parameters vary independently might be overly permissive, because combinations of these traits could be under stabilizing selection (we return to that shortly).

**Figure 4:**
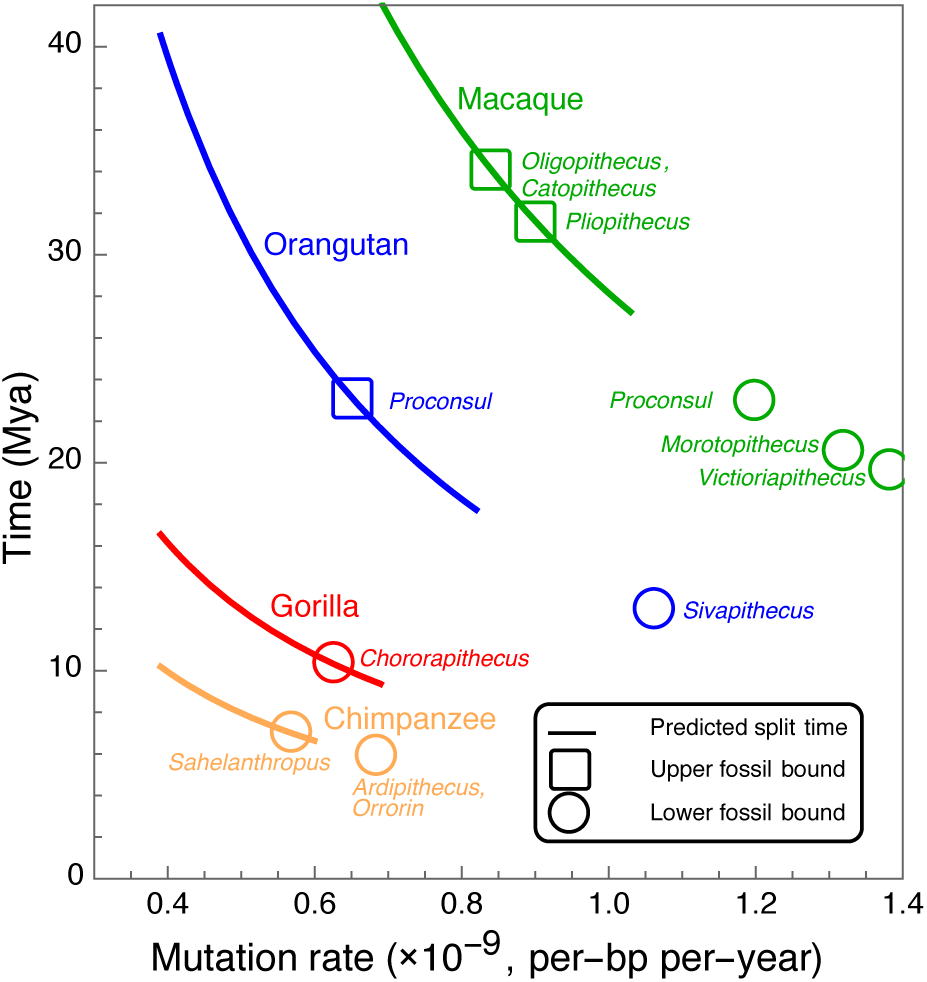
Estimated ranges of yearly mutation rates (x-axis) and species split times (y-axis) on the great ape phylogeny based on life history and spermatogenesis parameter estimates in extant species. The ranges of mutation rates on the human lineage following the split from chimpanzee (orange), gorilla (red), orangutan (blue) and macaque (green) are estimated as described in the text and in SI Section 5. To date the splits with orangutan and macaques, we follow Scally and Durbin (2012) in assuming an average ancestral coalescence time of 3 million years (39, 63). For the splits with chimpanzee and gorilla, we use estimates of the divergence after the split (38). Upper (squares) and lower (circles) bounds on spit times are based on the hypothesized phylogenetic positioning of fossils (cf. (62); SI Section 5).

Moving to broader phylogenetic contexts, we ask how our results suggesting that the average generation time has a moderate effect in hominines (also cf. (5, 24)) can be reconciled with molecular evolution studies showing that it is a major predictor of the rates of neutral substitutions in mammals (6, 10, 11). A possible explanation may arise from the relationship between generation time and the pre and post-pubertal number of germ cell divisions. For example, comparing mice, with an average generation time of ∼9 months, and humans, with an average generation time of ∼30 years, the estimated numbers are: 25 and 31 in females, 27 and 34 pre-puberty in males, and 35 and 390 during male spermatogenesis, respectively ((64) and Table S1). If we assume that more generally only the number of germ cell divisions during spermatogenesis increases vastly with the generation time (but see (26)) and that the number of germ-cell divisions determines the number of mutations, then several implications would follow. First, it would suggest that many of the mutations in great apes occur during spermatogenesis because of their exceptionally long generation times. It would further indicate that in species with short generation times, the contribution of mutations during spermatogenesis would be relatively small and that the classic model of a constant mutation rate per-generation may provide a reasonable approximation; or equivalently, that the yearly mutation rate would be approximately inversely proportional to the generation time (65).

To draw out these implications, we extrapolate our mutational model in hominines to a wider range of life history and spermatogenesis parameter values corresponding to a broader phylogenetic context (Figure 5). Such an extrapolation provides only a qualitative description of the dependency on the average generation time, as features and parameters of the mutational model likely vary over larger phylogenetic time scales. Notably, the mutation rate per cell division at different developmental stages might vary substantially among mammals ((60, 61) and Table S2). These reservations notwithstanding, we find that the average generation time dominates yearly mutation rates not only when the generation time is short but also when a wide range of generation times (and other life history traits) are considered (Figure 5A). The average generation time also affects the expected X-to-autosome ratio (with shorter generation times corresponding to lower *α* and a ratio closer to 1; (65)), though this effect is expected to be much more moderate and comparable with that of other life history traits (Figure 5B). Both of these predictions accord with the patterns found in mammals (6, 10, 11).

**Figure 5:**
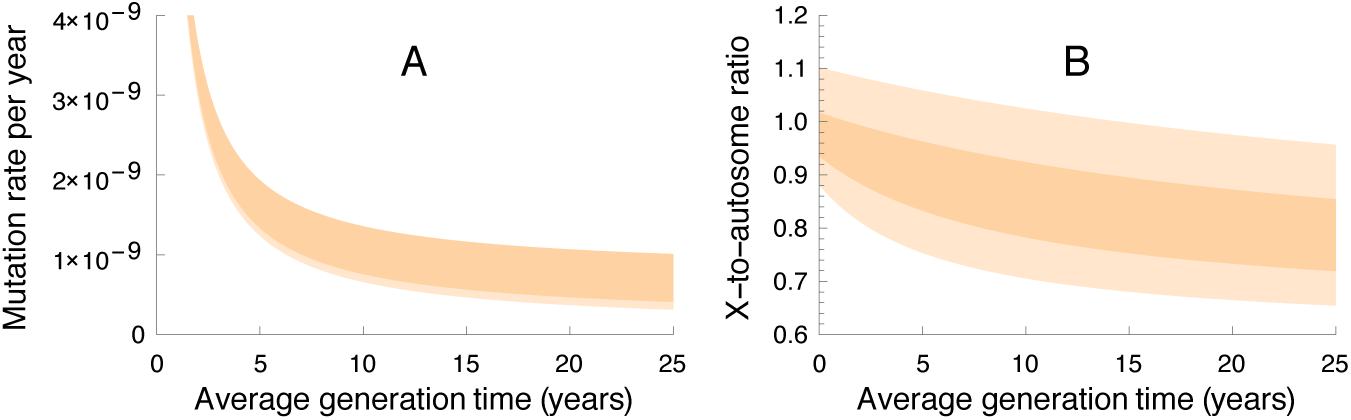
The effects of generation times on the molecular clock in a broader phylogenetic context. The yearly mutation rates (A) and X-to-autosome ratios (B) as a function of sex averaged generation time, based on our mutational model. The darker shade denotes narrower ranges of life history and spermatogenesis parameter values, corresponding to male-to-female ratios of generation times between 0.75 and 1.25, male ages of puberty and gestation times between 25-40% and 0-5% of their generation time, respectively (cf. (36) and Figure S1), and seminiferous epithelial cycles lengths between 6.5 and 16 days. The lighter shade denotes wider parameter ranges, corresponding to male-to-female ratios of generation times between 0.5 and 2, male ages of puberty and gestation times between 5-55% and 0-5% of their generation time, respectively, and seminiferous epithelial cycles lengths between 4 and 20 days.

It is interesting to contemplate how the relationship between generation time and germ cell divisions might affect and reflect selection on mutation rates. In so doing we set aside the question of selection on the mutation rate per cell division (cf. (60, 66)) and focus on selection on the number of germ cell divisions. Selection on mutation rates is expected to be stronger when mutation rates are higher, suggesting that it should be stronger in species with longer generation times (although species with longer generation times also tend to have lower effective population sizes; (60, 66)). Moreover, developmental constraints may limit the potential reduction in the number of pre-pubertal germ cell divisions, further suggesting a reduced efficacy of selection on mutation rates per generation in mammals with shorter generation times. In turn, selection on the number of divisions during spermatogenesis may be constrained by countervailing selection on the reproductive lifespan in males and on the amount of sperm they produce, e.g. imposed by sperm competition. Nonetheless, changes to the architecture of cell division in spermatogenesis may circumvent these countervailing constraints, allowing for effective selection for lower mutation rates in long-lived species (59). Albeit speculative, this reasoning lends support to the notion of stem cell turnover in spermatogenesis (59). Quantification of mutation rates in species with a wide range of generation times will shed light on these questions.

## Acknowledgments

We thank M. Przeworski for many helpful discussions throughout this work. We also thank P. Moorjani, M. Wyman, A. Scally and D. Reich for helpful discussions, P. Moorjani for sharing her unpublished results and M. Crist, D. Murphy, L. Hayward, G. Coop and M. Przeworski for comments on the manuscript.

## References

1. Kimura M (1984) The neutral theory of molecular evolution (Cambridge Univ. Press, London).

2. Kondrashov AS (2003) Direct estimates of human per nucleotide mutation rates at 20 loci causing Mendelian diseases. Hum. Mutat. 21(1):12–27.

3. Nachman MW & Crowell SL (2000) Estimate of the mutation rate per nucleotide in humans. Genetics 156(1):297–304.

4. Kong A, et al. (2012) Rate of de novo mutations and the importance of father’s age to disease risk. Nature 488(7412):471–475.

5. Segurel L, Wyman MJ, & Przeworski M (2014) Determinants of mutation rate variation in the human germline. Annu Rev Genomics Hum Genet 15:47–70.

6. Wu C-I & Li W-H (1985) Evidence for higher rates of nucleotide substitution in rodents than in man. Proceedings of the National Academy of Sciences 82(6):1741–1745.

7. Moorjani P, Amorim CEG, Arndt PF, & Przeworski M (2015) An unsteady molecular clock in primates. Manuscript in preperation.

8. Kim SH, Elango N, Warden C, Vigoda E, & Yi SV (2006) Heterogeneous genomic molecular clocks in primates. PLoS Genet 2(10):e163.

9. Patterson N, Richter DJ, Gnerre S, Lander ES, & Reich D (2006) Genetic evidence for complex speciation of humans and chimpanzees. Nature 441(7097):1103–1108.

10. Sayres MAW, Venditti C, Pagel M, & Makova KD (2011) Do variations in substitution rates and male mutation bias correlate with life-history traits? A study of 32 mammalian genomes. Evolution 65(10):2800–2815.

11. Ohta T (1993) An examination of the generation-time effect on molecular evolution. Proc. Natl. Acad. Sci. U. S. A. 90(22):10676–10680.

12. Sayres MAW & Makova KD (2011) Genome analyses substantiate male mutation bias in many species. Bioessays 33(12):938–945.

13. Presgraves DC & Yi SV (2009) Doubts about complex speciation between humans and chimpanzees. Trends Ecol. Evol. 24(10):533–540.

14. Michaelson JJ, et al. (2012) Whole-genome sequencing in autism identifies hot spots for de novo germline mutation. Cell 151(7):1431–1442.

15. Venn O, et al. (2014) Nonhuman genetics. Strong male bias drives germline mutation in chimpanzees. Science 344(6189):1272–1275.

16. Jiang YH, et al. (2013) Detection of clinically relevant genetic variants in autism spectrum disorder by whole-genome sequencing. Am. J. Hum. Genet. 93(2):249–263.

17. Penrose LS (1955) Parental age and mutation. Lancet 269(6885):312–313.

18. Crow JF (2000) The origins, patterns and implications of human spontaneous mutation. Nat Rev Genet 1(1):40–47.

19. Haldane JBS (1935) The rate of spontaneous mutation of a human gene. J. Genet. 31(3):317–326.

20. Haldane J (1946) The mutation rate of the gene for haemophilia, and its segregation ratios in males and females. Ann Eugen 13(1):262–271.

21. Pudney J (1995) Spermatogenesis in nonmammalian vertebrates. Microsc. Res. Tech. 32(6):459–497.

22. Franchi L, Mandl AM, & Zuckerman S (1962) The development of the ovary and the process of oogenesis. The ovary 1:1–88.

23. Nieuwkoop PD & Sutasurya LA (1979) Primordial germ cells in the chordates: embryogenesis and phylogenesis (Cambridge Univ. Press, London).

24. Gao Z, Wyman MJ, Sella G, & Przeworski M (2015) Interpreting the dependence of mutation rates on age and time. ArXiv e-prints. 1507:6890.

25. Johnson J, Canning J, Kaneko T, Pru JK, & Tilly JL (2004) Germline stem cells and follicular renewal in the postnatal mammalian ovary. Nature 428(6979):145–150.

26. Johnson J, et al. (2005) Oocyte generation in adult mammalian ovaries by putative germ cells in bone marrow and peripheral blood. Cell 122(2):303–315.

27. Ehmcke J, Wistuba J, & Schlatt S (2006) Spermatogonial stem cells: questions, models and perspectives. Hum. Reprod. Update 12(3):275–282.

28. Heller CG & Clermont Y (1964) Kinetics of the Germinal Epithelium in Man. Recent Prog. Horm. Res. 20:545–575.

29. Nielsen CT, et al. (1986) Onset of the Release of Spermatozia (Supermarche) in Boys in Relation to Age, Testicular Growth, Pubic Hair, and Height. J. Clin. Endocrinol. Metab. 62(3):532–535.

30. Fenner JN (2005) Cross-cultural estimation of the human generation interval for use in genetics-based population divergence studies. Am. J. Phys. Anthropol. 128(2):415–423.

31. Smithwick EB, Young LG, & Gould KG (1996) Duration of spermatogenesis and relative frequency of each stage in the seminiferous epithelial cycle of the chimpanzee. Tissue Cell 28(3):357–366.

32. Marson J, Meuris S, Cooper R, & Jouannet P (1991) Puberty in the male chimpanzee: progressive maturation of semen characteristics. Biol. Reprod. 44(3):448–455.

33. Langergraber KE, et al. (2012) Generation times in wild chimpanzees and gorillas suggest earlier divergence times in great ape and human evolution. Proc. Natl. Acad. Sci. U. S. A. 109(39):15716–15721.

34. Elango N, Thomas JW, & Soojin VY (2006) Variable molecular clocks in hominoids. Proc. Natl. Acad. Sci. U. S. A. 103(5):1370–1375.

35. Kingsley SR (1988) Physiological development of male orang-utans and gorillas. Orangutan Biology, ed Schwartz JH (Oxford Univ. Press, New York).

36. De Magalhães JP, Costa J, & Church GM (2007) An analysis of the relationship between metabolism, developmental schedules, and longevity using phylogenetic independent contrasts. J. Gerontol. A Biol. Sci. Med. Sci. 62(2):149–160.

37. Martin RD (2007) The evolution of human reproduction: a primatological perspective. Am. J. Phys. Anthropol. 134(S45):59–84.

38. Scally A, et al. (2012) Insights into hominid evolution from the gorilla genome sequence. Nature 483(7388):169–175.

39. Scally A & Durbin R (2012) Revising the human mutation rate: implications for understanding human evolution. Nat Rev Genet 13(10):745–753.

40. Gluckman PD & Hanson MA (2006) Changing times: the evolution of puberty. Mol. Cell. Endocrinol. 254-255:26–31.

41. Graves RR, Lupo AC, McCarthy RC, Wescott DJ, & Cunningham DL (2010) Just how strapping was KNM-WT 15000? J. Hum. Evol. 59(5):542–554.

42. Bauer VL & Aquadro CF (1997) Rates of DNA sequence evolution are not sex-biased in Drosophila melanogaster and D. simulans. Mol. Biol. Evol. 14(12):1252–1257.

43. Ellegren, H & Fridolfsson AK (1997) Male-driven evolution of DNA sequences in birds. Nat. Genet. 17(2):182–184.

44. Filatov, DA & Charlesworth D (2002) Substitution rates in the X-and Y-linked genes of the plants, Silene latifolia and S. dioica. Mol. Biol. Evol. 19(6):898–907.

45. Li WH, Yi S, & Makova K (2002) Male-driven evolution. Curr. Opin. Genet. Dev. 12(6):650–656.

46. Whittle CA & Johnston MO (2002) Male-driven evolution of mitochondrial and chloroplastidial DNA sequences in plants. Mol. Biol. Evol. 19(6):938–949.

47. Ellegren, H & Fridolfsson AK (2003) Sex-specific mutation rates in salmonid fish. J. Mol. Evol. 56(4):458–463.

48. Sandstedt SA & Tucker PK (2005) Male-driven evolution in closely related species of the mouse genus Mus. J. Mol. Evol. 61(1):138–144.

49. Berlin S, et al. (2006) Substitution rate heterogeneity and the male mutation bias. J. Mol. Evol. 62(2):226–233.

50. Bachtrog D (2008) Evidence for male-driven evolution in Drosophila. Mol. Biol. Evol. 25(4):617–619.

51. Harlid AB, Berlin S, Smith NG, Mosller AP, & Ellegren H (2003) Life history and the male mutation bias. Evolution 57(10):2398–2406.

52. Miyata T, Hayashida H, Kuma K, Mitsuyasu K, & Yasunaga T (1987) Male-driven molecular evolution: a model and nucleotide sequence analysis. Cold Spring Harb. Symp. Quant. Biol., pp 863–867.

53. Clutton-Brock TH & Isvaran K (2007) Sex differences in ageing in natural populations of vertebrates. Proc. Biol. Sci. 274(1629):3097–3104.

54. Dutheil JY, Munch K, Nam K, Mailund T, & Schierup M (2014) Strong selection in the human-chimpanzee ancestor links the X chromosome to speciation. bioRxiv:011601.

55. Charlesworth B (2001) The effect of life-history and mode of inheritance on neutral genetic variability. Genet. Res. 77(2):153–166.

56. Hwang DG & Green P (2004) Bayesian Markov chain Monte Carlo sequence analysis reveals varying neutral substitution patterns in mammalian evolution. Proc. Natl. Acad. Sci. U. S. A. 101(39):13994–14001.

57. Elango N, Lee J, Peng Z, Loh YH, & Yi SV (2009) Evolutionary rate variation in Old World monkeys. Biol. Lett. 5(3):405–408.

58. Forster P, et al. (2015) Elevated germline mutation rate in teenage fathers. Proc. Biol. Sci. 282(1803):20142898.

59. Scally A (2015) Mutation rates and the evolution of germline structure. Proc. R. Soc. B, Manuscript in preparation.

60. Lynch M (2010) Rate, molecular spectrum, and consequences of human mutation. Proc. Natl. Acad. Sci. U. S. A. 107(3):961–968.

61. Adewoye AB, Lindsay SJ, Dubrova YE, & Hurles ME (2015) The genome-wide effects of ionizing radiation on mutation induction in the mammalian germline. Nat Commun 6:6684.

62. Jensen Seaman MI & Hooper Boyd Ka (2013) Molecular Clocks: Determining the Age of the Human– Chimpanzee Divergence. eLS (John Wiley & Sons: Chichester) doi: 10.1002/9780470015902.a0020813.pub2.

63. Mailund T, Munch K, & Schierup MH (2014) Lineage sorting in apes. Annu. Rev. Genet. 48:519–535.

64. Drost JB & Lee WR (1995) Biological basis of germline mutation: comparisons of spontaneous germline mutation rates among drosophila, mouse, and human. Environ. Mol. Mutagen. 25(S2):48–64.

65. Li W-H, Ellsworth DL, Krushkal J, Chang BH-J, & Hewett-Emmett D (1996) Rates of nucleotide substitution in primates and rodents and the generation–time effect hypothesis. Mol. Phylogenet. Evol. 5(1):182–187.

66. Sung W, Ackerman MS, Miller SF, Doak TG, & Lynch M (2012) Drift-barrier hypothesis and mutation-rate evolution. Proc. Natl. Acad. Sci. U. S. A. 109(45):18488–18492.

